# Antibody Targeting of B7-H4 Enhances the Immune Response in Urothelial Carcinoma

**DOI:** 10.1101/730564

**Authors:** Joseph R. Podojil, Alexander P. Glaser, Dylan Baker, Elise T. Courtois, Damiano Fantini, Yanni Yu, Valerie Eaton, Santhosh Sivajothi, Paul Robson, Stephen D. Miller, Joshua J. Meeks

## Abstract

Locally advanced urothelial carcinoma has a poor survival despite a response to immunotherapy in approximately 25% of patients. To develop new therapies targeting other immune checkpoints, we identified increased expression of the T cell inhibitor protein, B7-H4 (VTCN1, B7S1, B7X), during tumor development of murine bladder cancer. Evaluation of B7-H4 in human bladder cancer by single-cell RNA-Seq and immune mass cytometry showed that B7-H4 is expressed by luminal tumors that are not normally responsive to PD-1 inhibitors. Additionally, overexpression B7-H4 was associated with significantly worse patient survival. In support of clinical translation, treatment of human monocyte and T cell co-cultures with a B7-H4 blocking antibody resulted in enhanced IFN-γ secretion by CD4^+^ and CD8^+^ T cell. While the study of B7-H4 in mouse cancer models has been difficult using tumor cell lines, our findings show that B7-H4 expression increased at 3 months after exposure to BBN on CD11b^+^ monocytes/macrophages and delivery of anti-B7-H4 antibody resulted in decreased tumor size and increased CD8 immune cell infiltration with increased CD8^+^/IFN-γ-producing T cells and a complimentary decrease in tumor infiltrating T regulatory cells (Tregs). Furthermore, treatment with a combination of anti-PD-1 and anti-B7-H4 antibodies resulted in a significant reduction in tumor stage. This data suggests that novel anti-B7-H4 antibodies maybe a viable target for bladder cancers unresponsive to PD-1 checkpoint inhibitors.

## Introduction

The anti-tumor immune response is dynamic and evasion of the immune system by the tumor is hallmark of cancer. Due to the expression of neoantigens driven by tumor-derived mutations, cells of the innate and adaptive immune system are capable of mounting vigorous anti-tumor responses. Yet, many tumors are able to evade this response by upregulating checkpoint inhibitors that suppress immune activation. While CTLA-4 and PD-L1 are the most well-described checkpoint proteins, B7-H4 (B7S1, B7x, VTCN1), a B7 family member expressed on antigen presenting cells and/or tumors, and can inhibit T cell-mediated inflammatory responses. The majority of work focused on elucidating the functional immunoregulatory role of B7-H4 has involved assessment of the ability of an agonistic B7-H4-immunoglobulin fusion protein (B7-H4Ig) in the treatment of autoimmune disease, and the presence of B7-H4 expression in human cancers [as reviewed in (1)]. B7-H4 functionally decreases inflammatory CD4^+^ T cell directly and indirectly via B7-H4-induced increases in both the number and function of regulatory CD4^+^ T cells (2–4). However, evidence for the ability of B7-H4 blockade to decrease tumor burden is lacking due to the limited number of mouse tumors that natively express B7-H4, likely secondary to non-orthotopic placement of ex-vivo derived cell lines. However, it was reported that genetic ablation of *B7-h4 (Vtcn1)* in a model of liver cancer was associated with increased CD8^+^ infiltration of the tumor and decreased exhaustion of CD8^+^ T cells (5).

In contrast to the majority of mouse tumor models, the expression level of B7-H4 protein in human ovarian cancer has been related to cancer types, cancer stage, the numbers of Tregs, and patient survival (6–8). Additionally, more than 90% of endometrial and breast cancers express B7-H4 protein (9), and B7-H4 expression in renal cell carcinoma (10–12), melanoma (13), breast (14), lung (15,16), gastric (17), colorectal (18), pancreatic (19), and prostate (20) cancer are associated with one or more clinicopathological factors, including tumor size, cancer stage, metastasis, progression, prognosis, survival, the intensity of tumor infiltrating T cells, the infiltration intensity of Tregs and tumor associated macrophages, disease recurrence or mortality. Therefore, targeting B7-H4 may be an alternative strategy to reinvigorate tumor-specific T cell responses.

Urothelial carcinoma is the fifth most common cancer in the US and has the second worst survival for patients with metastasis at only 5% at years (21). While systemic chemotherapy has been the standard of care for treatment of patients with metastatic urothelial carcinoma, in 2016 antibodies targeting immune checkpoint blockade (ICB), specifically PD1 and PDL1 were FDA approved (22). However, only 21% of patients with metastatic urothelial carcinoma refractory to chemotherapy respond to anti-PD1 antibodies (23). While the factors that determine clinical response are not completely known, features such as immune cell infiltration and high total mutation burden have been associated with an increased response (24). Not all studies have demonstrated that PDL1 expression is associated with improved survival following anti-PD-1 therapy, suggesting that multiple aspects of the regulation of immune responses to tumor remain unclear (25). Thus, most patients will progress and would benefit from additional therapies, targeting distinct, non-overlapping immune regulatory pathways.

## Results

### B7-H4 expression increases during bladder cancer development and is overexpressed in human bladder cancer

To investigate the interaction between bladder cancer initiation and progression we used the syngeneic, carcinogen-induced mouse model of bladder cancer derived from timed exposure to N-butyl-N-(4-hydroxybutyl)-nitrosamine (BBN). Our laboratory has previously validated the genomic credibility of this model that develops muscle-invasive bladder cancer (MIBC) between three and five months after initial exposure (37). Evaluation of the mutation signature most closely aligns with those caused by smoking, justifying the clinical relevance of BBN to bladder cancer, in which over 60% of cancers are attributed to smoking carcinogens (38). The BBN model of bladder cancer offers the distinct advantage that immune effector and regulatory processes can be assessed both in the bladder and peripheral immune organs from the initiation of tumorigenesis until the development of palpable tumors. Therefore, C57BL/6 (B6) mice were provided BBN and gene expression profiling was performed early (1, 2, and 4 weeks) after exposure to the carcinogen. We identified early expression of immune response-related genes that increased from low levels at one week to higher levels at two weeks of BBN exposure (Figure 1A). This included transcripts for CD8, CD4, and pathways associated with regulation of inflammation. For example, markers of T cell exhaustion (PD-L1, LAG-3 and Tim-3) and regulation (FoxP3 and Nrp-1) increased by 2 weeks, and B7-H4, a protein expressed by APCs that is known to inhibit T cell function via both direct and indirect mechanisms (4,39), was significantly increased by four weeks and corresponded to a decrease in CD8 and CD4 transcripts within the bladder (Figure 1A). To determine if B7-H4 could play a role in human bladder cancer, we evaluated the expression of B7-H4 in the cancer genome atlas for muscle invasive bladder cancer (MIBC). Patients with high expression of B7-H4 had significantly worse overall survival compared to those with lower expression (21.85 vs 34.03 months, p=0.03) (Figure 1B). Additionally, enriched B7-H4 expression positively correlated with B cells, iDCs, and macrophage transcripts, however B7-H4 expression did not correlate with T cells and IFN-γ levels present from MIBCs (29) (Supplemental Figure 1). Unlike other immune checkpoints (such as PD-L1, PD-L1, and CTLA-4), expression of B7-H4 does not appear to correlate with neoantigens, total mutation burden, APOBEC mutations or gene signatures (Supplemental Figure 2). We also found significant B7-H4 enrichment in bladder tumor subtypes that had relatively low expression of PD-L1, PD-1, and CTLA-4 including luminal, luminal papillary and luminal infiltrated tumors using the 2017 TCGA molecular subtyping classification (Figure 1C). Since expression of PD-L1 is directly associated with response to anti-PD-L1 and anti-PD-1 therapy, treatment of bladder cancer via blockade with a checkpoint inhibitor expressed in a different tumor subtype could lead to improved overall outcomes for patients. Thus, we were interested in the location of B7-H4 expression in bladder cancers, including localization of the tumor and tumor microenvironment. Using immunohistochemistry, we found expression of B7-H4 localized to immune cells resident in normal bladder, with increased expression in low and higher grade papillary tumors, respectively (Figure 1D, Right, middle panels). Yet, using the BBN mouse carcinogen model, B7-H4 expression co-localized with CD11b+ myeloid cells (Figure 1D, left) only. This suggests that expression of B7-H4 was likely localized to the myeloid lineage only and not expressed by the tumors in the BBN mouse model. To confirm this localization, we performed single-cell RNA-Seq of a stage Ta high grade bladder tumor (Figure 1E). *VTCN1* (B7H4) mRNA was detected on both urothelial cells (*EPCAM*+) and myeloid cells (*CD68*+) by scRNA sequencing. While *VTCN1* was expressed ubiquitously throughout the urothelial clusters, myeloid *VTCN1* was found on a subset of cells. Myeloid *VTCN1* shows scattered expression across the myeloid cluster without restriction to any one particular subcluster. To futher confirm the cellular localization of B7-H4, we performed imaging mass cytometry, that demonstrated localization of B7-H4 to CD68+ myeloid cells. Consistent with scRNA sequencing many CD68+ cells did not show expression of B7-H4 by imaging mass cytometry.

**Figure 1.**
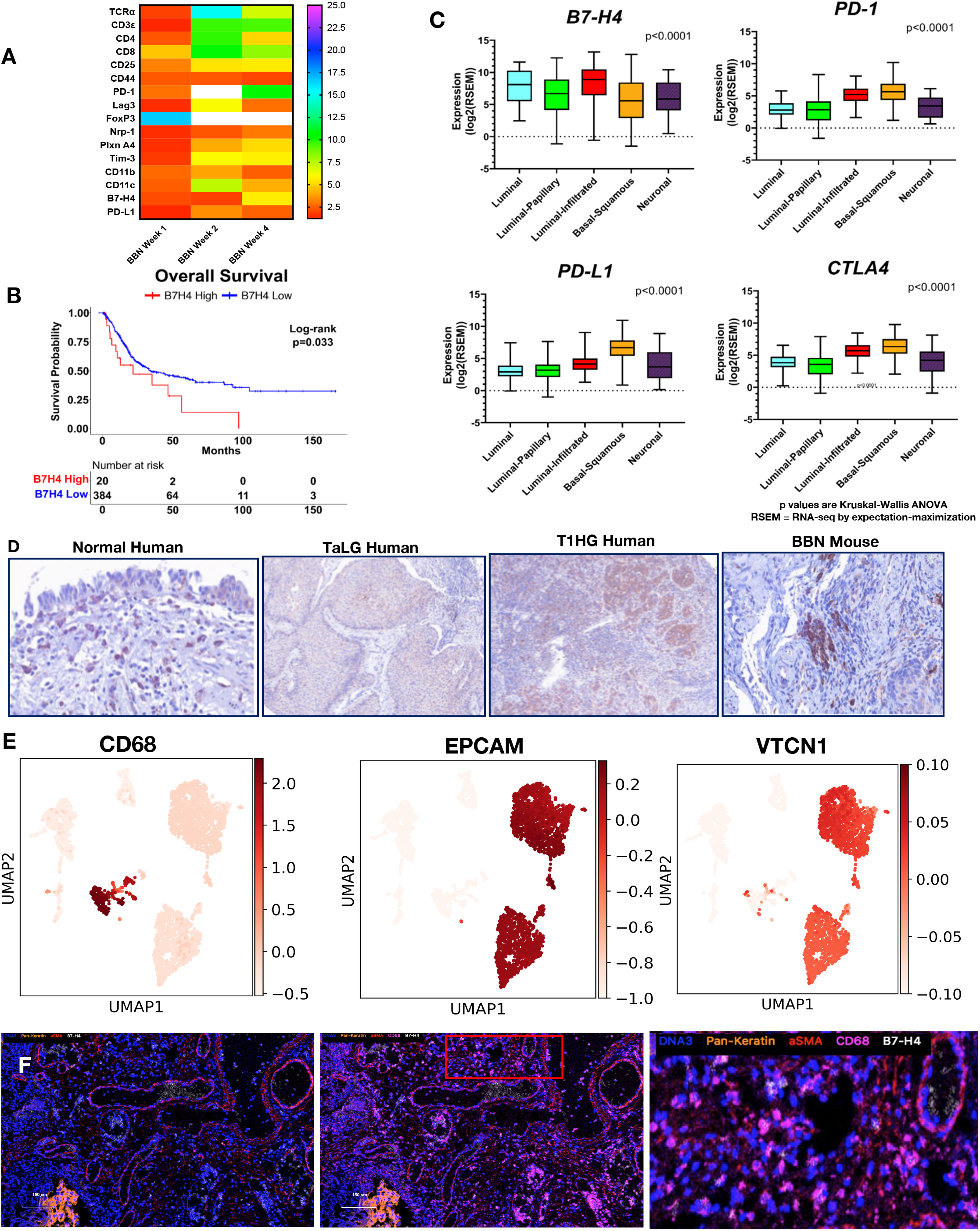
B7-H4 is expressed by bladder cancer and myeloid cells in human and murine bladder cancer. A) Heat map of mRNAs from immune regulatory genes at 1, 2 and 4 weeks of BBN exposure. B) Overall survival of the muscle invasive bladder cancers (MIBC) from the TCGA identifying worse survival for patients with high levels of VTCN1 (B7-H4). C) In contrast to tumors that express high levels of other immune regulatory proteins, like PD-L1, PD1 and CTLA4, B7-H4 is expressed in luminal and luminal infiltrated tumors, D) Immunohistochemistry of B7-H4 in normal human bladder, low grade non-invasive (stage TaLG), invasive (Stage T1HG) and BBN mouse bladder tumor demonstrating differences in cell localization. These localizations in the bladder were confirmed by single-cell RNA-seq of a low grade Ta tumor (E) showing high expression in luminal cells (+EPCAM) and scattered expression in myeloid cells (CD68). Immune mass cytometry (F) of a human bladder confirms localization of B7-H4 with CD68+ cells.

### Temporal expression of PD-L1 and B7-H4 and their respective ligands in BBN-induced bladder cancer

Mice given BBN in drinking water, develop carcinoma *in situ* (CIS) by 2-3 months, and mice present with tumors by 3 months and bulky tumors by 4 months (40). We next went on to validate the transcript data presented in Figure 1, by determining how the number of CD4^+^ T cells, CD8^+^ T cells, and monocytes/macrophages changed over time during BBN-induced bladder cancer development. Bladders (Figure 2) were collected at 0.5, 1, 2, 4, and 5 months after the initiation of BNN-induced bladder cancer. The results show a bi-phasic pattern in the number of CD4^+^ T cells, CD8^+^ T cells, and monocytes within the bladder over time (Figure 2A-C). Similar to our RNA-Seq results, we identified a peak in the numbers of pro-inflammatory T cells (CD4^+^ and CD8^+^) at 1 month, but a decline in cell numbers by 2 months. The numbers of bladder infiltrating CD4^+^ and CD8^+^ cells did not increase until 3 months concomitant with the development of carcinoma. During BBN exposure, the gradual increase in CD4^+^ and CD8^+^ T cells was associated with increased expression of the exhaustion marker, PD-1 (Figure 2D-E). The numbers of CD4^+^ T cells expressing the neuropilin-1/semaphorin3a (Nrp-1/Sema3a), B7-H4 receptor (39) was increased at 1 and 5 months (Figure 2G) while B7-H4 receptor-positive CD8^+^ T cells increased at 0.5 and 1 months (Figure 2H). Monocytes (CD11b^+^/Ly6C^+^/Ly6G^−^) increased sharply at 1 month and subsequently decreased concordant with the decrease in T cell numbers. The infiltrating monocytes expressed PD-L1 at 1 month with a second wave of increased expression at 5 months (Figure 2F). There was a small peak of B7-H4^+^ monocytes at one month, followed by a decline, and a second peak concordant with tumor development and immune cell infiltration of bladder tumors at 5 months (Figure 2I). Additionally, the bi-phasic pattern of PD-L1 and B7-H4 expression correlated with both the numbers of CD4^+^ Treg (Figure 2J) and effector CD4^+^ T and CD8^+^ T cells (Figure 2K and 2L). These findings confirm B7-H4 expression by bladder-infiltrating monocytes concordant with CD4^+^ and CD8^+^ TILs suggestive of induction of an anti-inflammatory regulatory mechanism. Based on these results, we investigated the blockade of B7-H4 as a potential therapy to increase T cell responses by blocking B7-H4-mediated T cell regulation.

**Figure 2.**
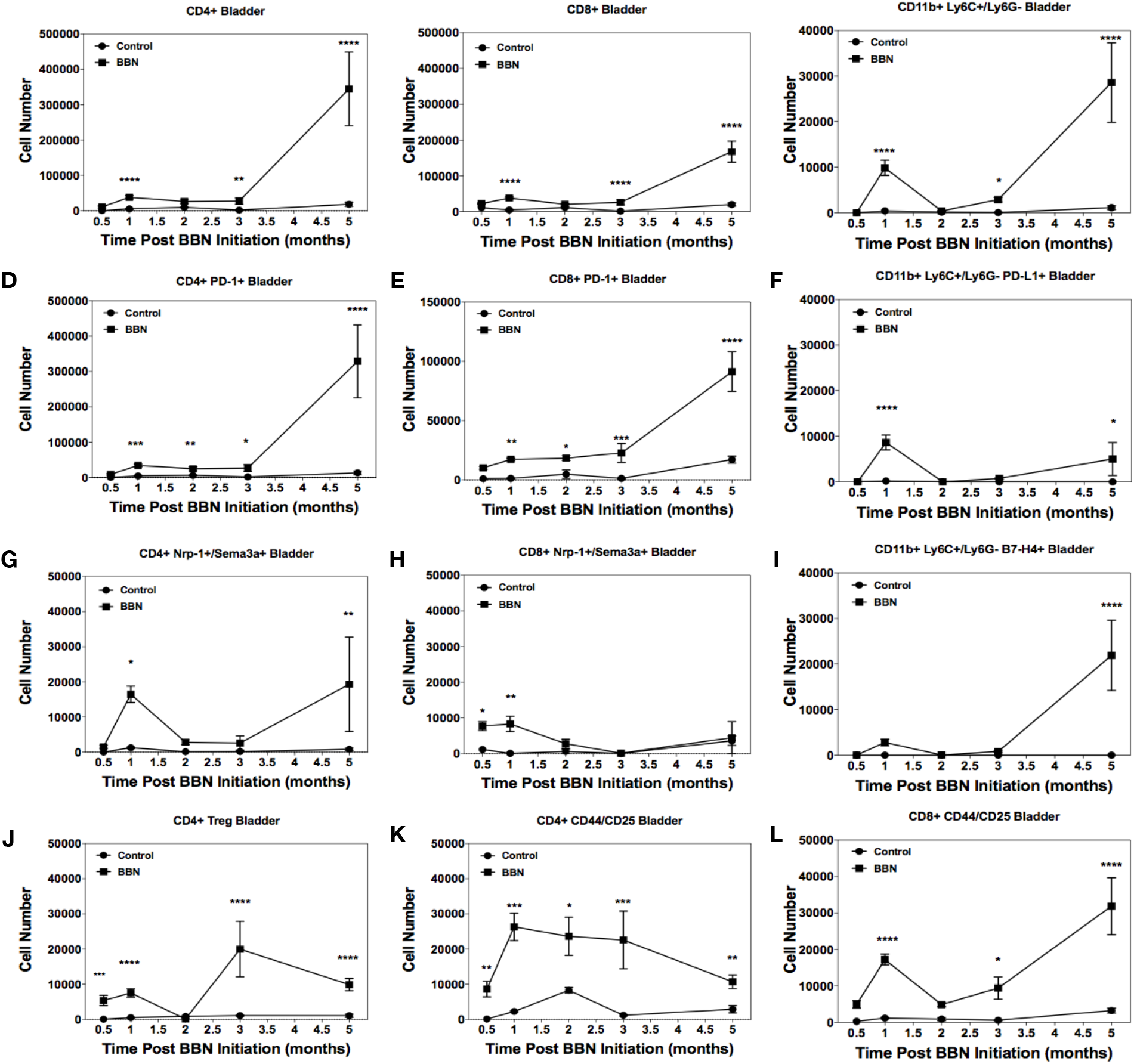
Bi-phasic pattern of immune bladder immune infiltration during BBN-induced cancer. Male C57BL/6 mice (n=10 mice per group for each time point) were supplied either sterile water or sterile water containing 0.05% BBN *ad libitum*. On the indicated time points, 10 representative naïve control and BBN treated mice were taken for analysis. The bladders were harvested and 2 random bladders each pooled to generate a total of 5 analytical samples for flow cytometric analysis. The cells samples were gated as follows; for T cells (singlets, cells, live/CD45^hi^, CD3^+^/CD4+ or CD3^+^/CD8^+^ into flow plots for specified T cells markers), and for monocytes (singlets, cells, live/CD45^hi^, CD11b^+^, Ly6C^+^/Ly6G^−^ into flow plots for specified monocyte markers). The data are presented as the mean number of each specified cell population per sample ± S.E.M. Asterisks indicate a statistically significant difference in the number of cells present within the bladder of mice receiving BBN as compared to Control naïve mice analyzed at each time point - *p<0.05, **p< 0.01, ***p< 0.001, ****p<0.0001, respectively.

### Anti-B7-H4 blocks the immunoregulatory function of B7-H4Ig and increases mouse and human CD4^+^ and CD8^+^ T cell function

To functionally demonstrate that anti-B7-H4 can inhibit the regulatory function of B7-H4, we tested the ability of anti-B7-H4 to block the immunoregulatory function of human B7-H4Ig. SJL/J lymph node cells (5×10^5^ cells/well) were labeled with CFSE, and stimulated with anti-CD3 (1μg/ml) plus control Ig, human B7-H4Ig, and/or anti-B7-H4 (10μg/ml) as indicated (Figure 3A). On Day 3 of culture, the cells were collected and the level of CD4^+^ T cell CFSE dilution assessed via flow cytometry. While anti-B7-H4 alone did not alter the extent of CD4^+^ T cell proliferation, the addition of anti-B7-H4 inhibited hB7-H4Ig-induced suppression of CD4^+^ T cell proliferation. As a secondary confirmation of the CFSE findings, replicate wells were pulsed with ^3^H-TdR on Day 1 and cultures harvested on Day 3. The level of ^3^H-TdR incorporation (counts per minute, CPM) confirmed that anti-B7-H4 blocked the inhibitory function of human B7-H4Ig (Figure 3A). As a further confirmation of the ability of anti-B7-H4 to block hB7-H4Ig function, SJL/J mice were primed with PLP_139-151_/CFA, and treated 3 times per week for 2 weeks beginning at the time of priming (Day 0) with control Ig, hB7-H4Ig, or anti-B7-H4 as indicated. Mice were observed for paralytic disease symptoms. As we previously reported (4), hB7-H4Ig treatment significantly decreased disease severity (Figure 3B). While anti-B7-H4 treatment alone did not modulate the level of disease severity, treatment with anti-B7H4 blocked the hB7-H4Ig-induced decrease in disease severity (Figure 3B). In addition, anti-B7H4 blocked the hB7-H4Ig-induced decrease in *in vivo* CD4^+^ T cell function as measured by delayed-type hypersensitivity (DTH) responses to ear challenge with PLG_139-151_. Taken together, these findings show that anti-B7-H4 blocks hB7-H4Ig-induced regulatory function on mouse T cell responses both *in vitro* and *in vivo*.

**Figure 3.**
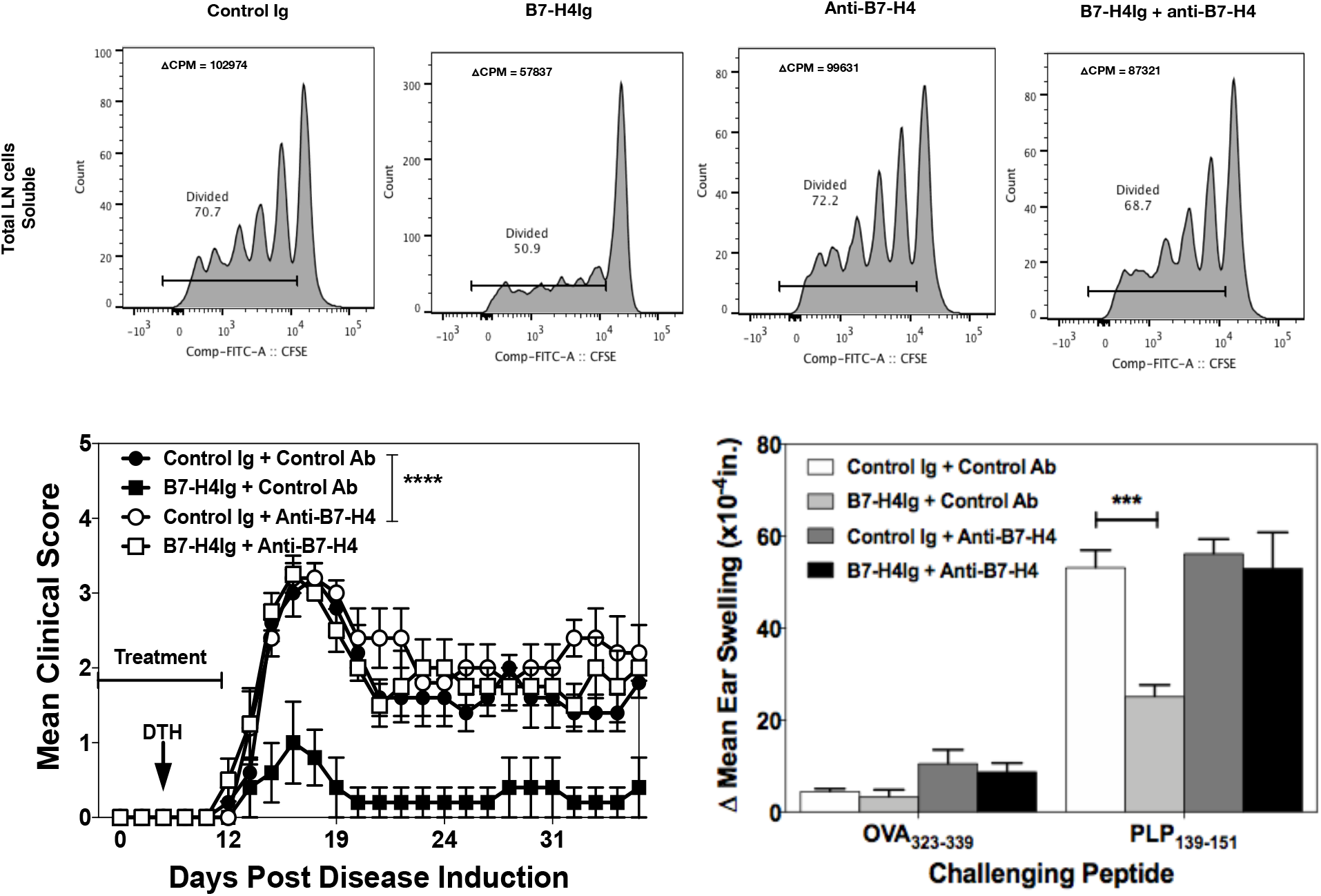
Anti-B7-H4 blocks the immunosuppressive function of B7-H4Ig. Total LN cells were collected from naïve SJL/J mice, and cells were labeled with CFSE. CFSE-labeled lymph node cells (5×10^5^ cells/well) were cultured in the presence of anti-CD3 (1μg/ml), plus Control Ig, hB7-H4Ig, anti-B7-H4, or hB7-H4Ig plus anti-B7-H4 (5μg/ml). Replicate wells were pulsed with 1μCi of tritiated thymidine at 24 h, and the cultures were harvested on day +5. For the assessment of T cells proliferation, the percentage of live CD4+ T cells that diluted CFSE were gated. The level of cellular proliferation is also presented on each flow histogram as determined by tritiated thymidine incorporation (Δ CPM = CPM with anti-CD3 – CPM with PBS) (A). SJL/J mice (n=10/group) were primed with PLP_139-151_/CFA and treated with species and isotype-matched Control Ig, hB7-H4Ig, anti-B7-H4, or hB7-H4Ig plus anti-B7-H4 (100μg/dose; 3x/wk; 2wks), and mice were followed for disease severity. The data is presented as the mean clinical score ± S.E.M. (B). On day +9 of the disease course the PLP_139-151_ *in vivo* DTH responses were assessed. The data are presented as the mean ear swelling ± S.E.M. for both control antigen challenge (OVA_323-339_) and specific antigen challenge (PLP_139-151_) (C). One representative experiment of three is presented. Asterisks (***) indicates a statistically significant decrease in the PLP_139-151_-induced clinical EAE or DTH response in mice B7-H4Ig treated mice in comparison to mice treated with Control Ig, p < 0.001 respectively.

To determine if anti-B7-H4 could block B7-H4-induced regulation of human T cells, CD14^+^ monocytes from healthy donors were cultured for 3 days with IL-10, IL-6, and IL-10 plus IL-6 (10ng/ml). Adherent cells were collected and analyzed via flow cytometry. Upregulated expression of B7-H4, CD80, CD86, PD-L1, and PD-L2 expression was identified following stimulation with IL-10 and/or IL-6 (Figure 4A). Therefore, IL-10 + IL-6 stimulated CD14^+^ APCs were co-cultured with autologous CD4^+^ and CD8^+^ T cells in anti-CD3-coated wells + control Ig or anti-B7-H4 and proliferation and IFN-γ production measured on 3 d thereafter. While addition of anti-B7-H4 did not significantly enhance proliferation of CD4^+^ (Figure 4B) or CD8^+^ T cells (Figure 4D), anti-B7-H4 induced a significant increase in production of IFN-γ by both CD4^+^ and CD8^+^ T cells (Figure 4C and 4E). Thus, anti-B7-H4 blockade increased the function of both human CD4^+^ and CD8^+^ T cells stimulated with B7-H4 expressing monocytes.

**Figure 4.**
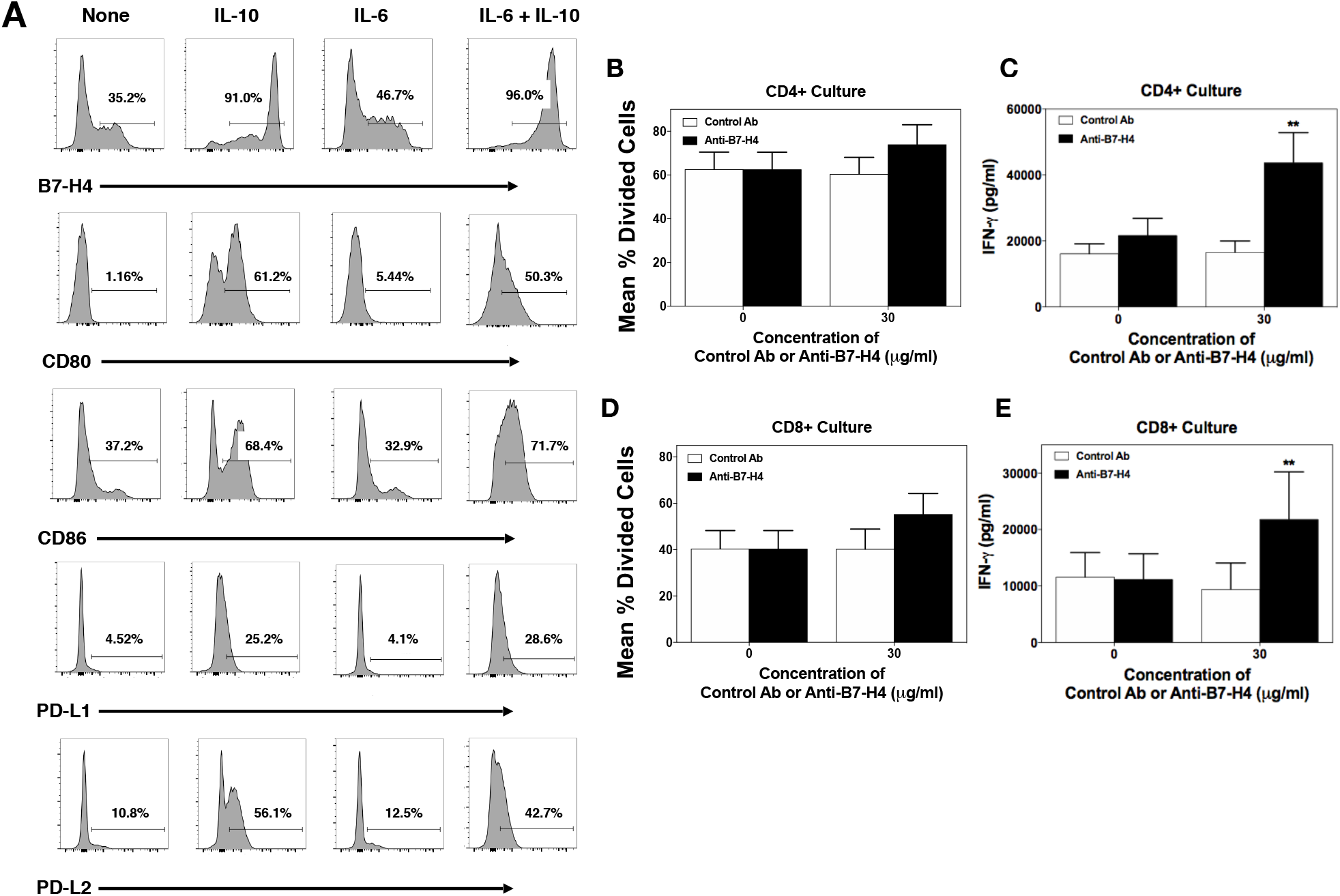
Anti-B7-H4 treatment increase IFN-γ secretion by T cell cultured in the presence of B7-H4+ monocytes. CD14^+^ monocytes were sort purified from healthy donor PBMCs (n=6) and cells were cultured in the presence of medium alone, IL-10, IL-6, or IL-10 plus IL-6 (20ng/ml) for 3 days. The percent of resultant monocytes expressing B7-H4, CD80, CD86, PD-L1, and PD-L2 was assessed (A). Sort purified autologous CFSE-labeled CD4^+^ T cells (B, C) and CD8+ T cells (D, E) were co-cultured with IL-10 plus IL-6 conditioned monocytes and anti-CD3 for 3 days, and the level of proliferation was assessed via flow cytometry (B, D) and the level of secreted IFN-γ measured (C, E). Proliferation data is presented as the mean percent divided cells and concentration of IFN-γ (pg/ml) present with the culture ± S.E.M. Asterisks (**) indicates a statistically significant difference as compared to the Control Ab treated cultures, p<0.01, respectively.

### Anti-B7-H4 treatment increases CD8^+^ T cell infiltration into the bladder during BBN-induced bladder cancer and IFN-γ production by splenic CD8^+^ T cells

After confirmation of the functional activity of our B7-H4 antibody, we tested its activity in the BBN carcinogen model of bladder cancer. Mice received BBN containing water for four months and received treatment with a species and isotype matched Control Ab, anti-PD-1, or anti-B7-H4 for once weekly for four total treatments. After treatment, animals were placed on normal drinking water for 30 days followed by tumor analysis (Figure 5A). Anti-B7-H4 treatment led to a decrease in the number of advanced tumors (70% compared to 50% stage 3 or greater), and a significant increase in the number of CD8^+^ bladder-infiltrating T cells (Figure 5B, C). In contrast, anti-B7-H4 treatment decreased the number of FoxP3+ Tregs (Fig 5B,D), indicating that anti-B7-H4 treatment had shifted the CD8^+^ T cell to Treg ratio in favor of an anti-tumor response. While treatment of mice with anti-PD-1 or anti-B7-H4 did not alter the number of total splenic CD45hi cells, CD4^+^ T cells, or CD8^+^ T cells (Figure 5E), both anti-PD-1 and anti-B7-H4 treatments significantly increased the number of CD8^+^/IFN-γ^+^ cells present within the spleen compared to control IgG-treated mice (Figure 5F). Additionally, total splenocytes from the treatment groups were activated *ex vivo* in the presence of anti-CD3 for 3 days to assess levels of cytokine secretion. Significantly, splenocytes from anti-B7-H4-treated mice secreted significantly higher levels of IFN-γ in response to anti-CD3 stimulation than did T cells from control IgG or anti-PD-1 treated mice (Figure 5G). Taken together, these findings indicate that anti-B7-H4 treatment of BBN mice induced an increase in CD8^+^ T cell infiltration into tumors within the bladder and increased the level of IFN-γ secreted per splenic CD8^+^ T cell.

**Figure 5.**
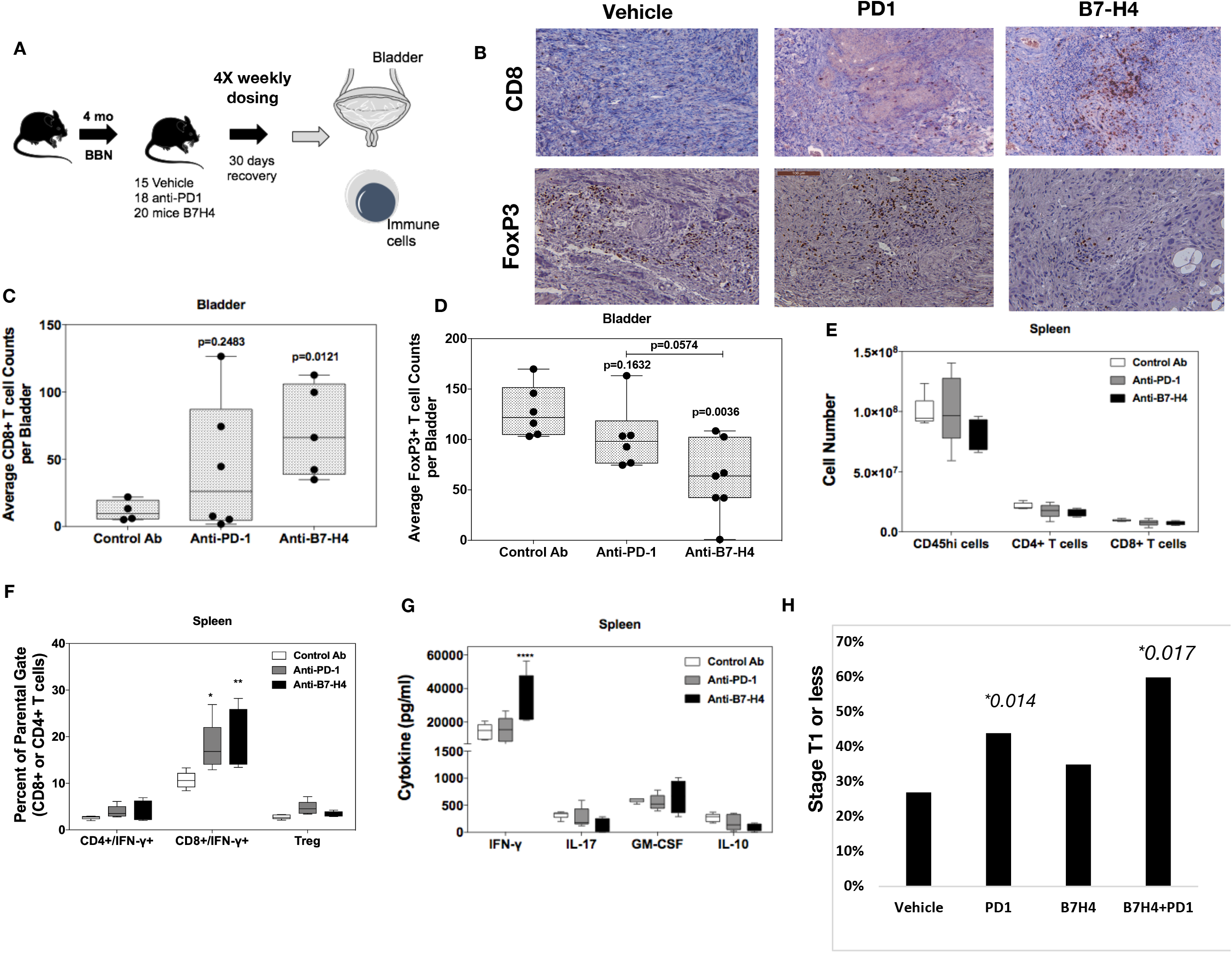
Anti-B7-h4 treatment increases CD8+ T cell function. (A) Treatment schema of mice for comparison of anti-tumor response of IgG, anti-PD1 and anti-B7-H4 antibodies. (B) Representative immunohistochemistry sections of BBN tumors for anti-CD8 (top) and anti-FoxP (bottom) staining. All images are at low-power (100X). Quantification of the average number of CD8+(C) and FoxP3 (D) immune cells per low power field. The number (E) and percentage (F) of singlet/live/CD45hi cell that were CD4^+^/IFN-γ^+^, CD8^+^/IFN-γ^+^ or CD4^+^/CD25^+^/FoxP3^+^ present within the spleens was assessed. The data are presented as the mean cell number or percentage of cells ± S.E.M. Total splenocytes were cultured in the presence of anti-CD3 (1 μg/ml) for 3 days and the level of secreted cytokine assessed. The data are presented as the mean pg/ml of secreted cytokine (G). Asterisks (*, **, ****) indicates a statistically significant difference as compared to the Control Ab treated mice, p < 0.05, 0.01, 0.001 respectively. H) BBN mice treated with anti-PD1 and anti-B7H4 had lower rates of T2 or higher (detrusor invasion and greater) cancer compared to anti-PD1 or anti-B7-H4 alone.

We evaluated the gene expression profile of anti-B7H4 treated mice and found a re-establishment of normal gene expression associated with urothelium differentiation. These included increased expression of bladder muscle/detrusor-related pathways and urothelial differentiation (uroplakin 3b) and decreased expression of cell-cycle progression, mitosis, and cell cycle checkpoints (Supplemental Figure 3). Most prominent, response to B7-H4 was associated with decreased expression of oncogenes Hoxb9, H19 and Egln3. We also observed metabolic changes suggestive of aerobic pathways including Car2 (carbonic anhydrase), adh1 (alcohol dehydrogenase), and Pck1. Lastly, we showed that anti-B7H4 antibody therapy synergized with anti-PD1 therapy, the current standard of care for metastatic, and chemotherapy-resistant cancer, leading to a significant reduction in advanced tumors (40% increase in stage I tumors: 60% in combination treated mice compared to 20% in vehicle treated mice (Figure 5H) (p=0.017).

## Discussion

Bladder cancer, similar to other solid tumors attributed to carcinogen exposure including lung cancer and melanoma, is driven by a high total mutation burden at approximately 7 mutations/MB (41). Systemic immunotherapy has changed the survival of patients with bladder cancer for those unresponsive to cisplatin (CIS) chemotherapy, or unable to received chemotherapy with a median progression free survival of 10.3 months (95% confidence interval [CI], 8.0 to 11.8) compared with 7.4 months (95% CI, 6.1 to 8.3) in those treated with chemotherapy (hazard ratio for death, 0.73; 95% CI, 0.59 to 0.91; P=0.002) (23). Recent application of anti-PD1 and anti-PDL1 therapy to the neoadjuvant setting have shown promising results to decrease pathologic stage and potentially improve recurrence free survival (42). Yet, less than 20% of patients treated with checkpoint immunotherapy have a clinical response and even in those with MIBC, choosing the right systemic therapy prior to cystectomy remains critical to limit toxicity prior to resection. Until the early reports of Phase III trials of Atezolizumab and Pembrolizumab in the first-line metastatic setting, the value to PD-L1 testing was unclear. Yet, inferior reports of response to PD-L1 low or non-expressing tumors in May of 2018 resulted in label changes for both Atezolizumab and Pembrolizumab confirming the critical association of PD-L1 immunohistochemical marker activity with response to each therapy. Thus, patients that are PD-L1 “negative” have a low likelihood of response and are unlikely to benefit from systemic therapy targeting PD-1/PD-L1. Due to the limitations of the current landscape of PD-1/PDL-1 therapy for bladder cancer, investigation of other immune checkpoints is a necessity for those patients unlikely to respond to PD-1/PD-L1. In our evaluation of MIBC, we found that B7-H4 is expressed in higher levels in luminal and luminal papillary tumors. This distinction separates B7-H4 from CTLA-4 and PD-L1 which are expressed at higher levels in basal cancers. While tumor subtyping has been performed in small series of patients treated with anti-PD-1/PD-L1, luminal and luminal papillary tumors are the poorest responder (22,43)

To identify potential immune regulatory mechanisms involved in the immune-editing of bladder cancer, we profiled the immune landscape of the mouse BBN bladder cancer model from 2 to 20 weeks. Our group has previously validated the genomic alterations that develop in this model (37). To our knowledge, we are the first to describe the immune activation that is bi-modal after BBN treatment. The present data show that there is an acute inflammatory phase that starts within two weeks, but then decreases between four and twelve weeks. While we observed invasion of the bladder by lymphocytes between two and four weeks post initiation of BBN treatment, expression of regulatory monocyte and T cell populations potentially decrease the effector CD4^+^ and CD8^+^ T cells until the tumor develops at three months. With the development of CIS and invasive bladder cancer, the number tumor infiltrating lymphocytes increases, but these cells express markers of exhaustion (PD-L1) indicating they are likely not effective in controlling tumor cell growth. Upregulation of B7-H4 expression was observed in monocytes in the BBN-treated mice, preceding both the decrease in CD8^+^ cells and increase in Tregs. This led to further functional studies of B7-H4 in the BBN bladder cancer model.

Across solid tumors, expression of B7-H4 is associated with worse survival (6–8). High expression of B7-H4 has been described in ovarian, endometrial, gastric, renal cell, melanoma, breast, lung, colorectal, pancreatic, and prostate cancer. Mechanistic investigation of B7-H4 in liver cancer identified B7-H4 expression on tumor cells and antigen presenting cells of the liver that were coexpressed with PD-1 and Tim-3. Genetic targeting of B7-H4 resulted in increased proliferation of CD8^+^ T cells with decreased levels of exhaustion transcription factors (5). In the current study, we confirmed increased CD8^+^ TIL activation induced by treatment with anti-B7-H4 antibody as a mechanism of immune activation. Additionally, we found lower numbers of FoxP3^+^ Tregs that can decrease the anti-tumor immune response.

In addition to effects on Tregs, receptor ligation by the immune modulatory molecules, such as B7-H4, have been suggested to directly alter the phenotype of CD4^+^ T cells (44,45). Co-culture of CD4^+^ T cells with IL-10/TGF-β treated macrophages expressing B7-H4 decreased CD4^+^ T cell proliferation and increased the number of FoxP3^+^ CD4^+^ T cells (46). Published data show that hB7-H4Ig binds to CD4^+^ T cells in an activation-dependent manner, indicating that B7-H4 may function as a co-inhibitory (negative co-stimulatory) molecule for CD4^+^ T cells (4,44,47). The present data show that B7-H4 also directly interacts with a functional B7-H4 receptor on CD8^+^ T cells, as the addition of anti-B7-H4 to monocyte/CD8^+^ T cell co-cultures induced a significant increase in the level of secreted IFN-γ (Figure 4E). This is supported by our previous findings showing that hB7-H4Ig can down-regulate IL-17 and IFN-γ production of mouse T cells in the absence of Tregs (4). Further, our *in vivo* data show that treatment of C57BL/6 mice with BBN-induced bladder cancer have a significant increase in the number of CD8^+^ T cells and a significant decrease in the number of Tregs present within the bladder (Figure 5C and D). This finding correlated with an increase in the percentage of IFN-γ^+^/CD8^+^ T cells present within the spleen and a significant decrease in tumor stage as compare to Control Ab treated mice. While the present data have shown the functionality of anti-B7-H4 treatment in the presence of monocyte expressed B7-H4, B7-H4 expression by tumor cells is also a putative mechanism by which these cells evade anti-tumor immune responses. In the majority of breast and ovarian cancers, B7-H4 mRNA is expressed at approximately 2-fold or greater than the level expressed within normal tissue (48), and B7-H4 protein is present in half of early stage and two-thirds of late stage ovarian tumors (49). Additionally, tumor tissues from breast, uterus, ovary, colon, and pancreas showed a statistically significant increase in the percentage of cells expressing B7-H4 (50). In the 4T1 metastatic breast cancer model, transfer of tumor cells into B7-H4^−/−^ vs. wildtype mice resulted in fewer lung nodules, enhanced survival, and decreased tumor infiltration of immunosuppressive cells (51). Based on these findings expression of B7-H4 within the tumor microenvironment, either by the tumor cells and/or by infiltrating monocytes, is hypothesized to promote immune evasion. In support of this hypothesis, we showed that stimulation of the B7-H4 receptor complex via hB7-H4Ig treatment of mice during EAE increased the number and function of Tregs (4).

Our investigation identified B7-H4 as a negative regulator of T cell activity in the bladder during cancer development. The present data show that treatment of C57BL/6 male mice receiving BBN containing water with anti-B7H4 increases the number of CD8+ T cells and decreases the number of Tregs present within the bladder. The anti-B7-H4 treatment-induced skewing of the CD8^+^ T cell to Treg ratio within the bladder correlated with a decrease in the bladder cancer stage score. Additionally, co-treatment of mice with both anti-B7-H4 and anti-PD1 further decreased bladder cancer stage score, and such combination therapy may be considered a target for future therapeutic trials in patients with bladder cancer.

## Materials and Methods

### Tumor Preparation for Single-Cell RNA-Seq

Tumor samples were obtained prospectively after IRB approval at Northwestern (STU00088853). Tumor specimen was minced and enzymatically dissociated DMEM supplemented with Liberase TM (0.0625 mg/ml) and DNase I (Sigma, D5025, 0.2 mg/mL) for 30 minutes. Every 10 minutes specimen was gently pipetted and enzyme mix was exchanged for freshly made enzyme mix. After dissociation tissue was spun down at 1300 RPM for 7 minutes and filtered to through a 100 um filter to yield a single cell suspension. Cells were spun down, resuspended in PBS supplemented with 0.5% BSA and 2 mmol/L EDTA and stained with PI (BD) and Calcein Violet (Invitrogen). Viable cells were sorted using BD FACS Aria Fusion instrument. Sorted cells were washed and resuspended in PBS containing 0.04% BSA. Cells were counted on Countess II automated cell counter (Thermo Fisher) 12,000 cells were loaded per lane onto a 10X Chromium microfluidic chip. Single-cell capture, barcoding, and library preparation were performed using the 10X Chromium version 2 chemistry according to the manufacturer’s protocol (#CG00052). cDNA and libraries were checked for quality on Agilent 4200 Tapestation and quantified by KAPA qPCR before sequencing on a single lane of a HiSeq4000 (Illumina) to an average depth of 50,000 reads per cell.

### Single-Cell Data Processing

The Cell Ranger pipeline (v1.2, 10X Genomics) was used to convert Illumina base call files to FASTQ files, align FASTQs to the GRCH38 reference (v3.0.0, 10X Genomics) for human samples to produce a digital gene-cell counts matrix. The resultant gene-cell matrix was filtered to remove cells with fewer than 500 transcripts and genes with fewer than two counts in two cells. The genecell matrices were then normalized such that the number of unique molecular identifiers (UMI) in each cell is equal to the median UMI count across the data set and log transformed. Expression at 1,000 highly variable genes in each data set, selected as the genes with the highest dispersion, was used to reduce the dimensionality of the data sets to three dimensions using Uniform Manifold Approximation and Projection (UMAP) and cells were clustered using Leiden based clustering in the UMAP space. Genes of interest were plotted in UMAP space using adjusted values based on Markov Affinity-based Graph Imputation (MAGIC) of the raw gene-cell counts matrix.

### Multiplexed Imaging by IMC

Formalin-fixed paraffin-embedded human Ta NMIBC tissues were cut into 5-μm sections and mounted on slides. Slides were incubated for 15 minutes at 55°C in a dry oven, deparaffinized in fresh histoclear, and rehydrated through a series of graded alcohols. Antigen retrieval was performed in a decloaking chamber (BioSB TintoRetriever) for 15 minutes at 95°C in Citrate Buffer. After blocking in buffer containing 3% BSA, slides were incubated overnight at 4°C with a cocktail of metal-conjugated IMC-validated primary antibodies. The following day, slides were washed twice in Dulbecco’s Phosphate-Buffered Saline and counterstained with iridium intercalator (0.25 [μmol/L) for 5 minutes at room temperature, to visualize the DNA. After a final wash in ddH20, the slides were air-dried for 20 minutes. The slides were then loaded on the Fluidigm Hyperion imaging mass cytometer. Regions of interest were selected using the acquisition software and ablated by the Hyperion. The resulting images were exported as 16-bit.tiff files using the Fluidigm MCDViewer software and analyzed using the open source Histocat++ toolbox

### Mice and tumor staging

Mice were cared for in a fashion supervised by the ACUC at Northwestern University and all research was approved in an animal research protocol. Male C57BL/6 mice at least 6 weeks old received N-butyl-N-(4-hydroxybutyl)-nitrosamine (BBN) at a dose of 0.05% in drinking water *ad libitum*. After 20 weeks of exposure, mice were randomized to therapy which included treatment with IgG (200 μg/kg, BioXcell), anti-PD1 (200 μg/kg, BioXcell), and/or anti-B7-H4 (200 μ.g/mouse, from Amplimmune). Mice were injected with four weekly doses. At the end of treatment, animals were euthanized by CO_2_ with secondary cervical dislocation.

Bladders were removed and divided in the sagittal plane, and either fixed in 10% formalin or snap-frozen in liquid nitrogen. Pathology was staged using human AJCC staging of degree of bladder wall invasion.

### RNA extraction, Library Creation and RNA-Seq

Mouse tumors were ground using liquid nitrogen and a mortar and pestle, and then RNA was extracted using Trizol Reagent (Thermo Fisher Scientific) following manufacturer’s instructions. RNA quality and concentration were assessed by NanoDrop and then Qubit Fluorometric Quantitation (Thermo Fisher Scientific). Library preparation and RNAseq were performed at the NuSeq core facility (Northwestern University) using an Illumina HiSeq 2000.

### Immunohistochemistry

Bladders were fixed in 10% formalin and embedded in paraffin. 4 micron thick sections were used for IHC staining for CD8 (1:1000, #MA5-13263, Invitrogen) and FoxP3 (1:400, #12653, Cell Signaling). IHC was performed using a Dako Autostainer Plus instrument (Dako, CO, USA), and anti-rabbit Dako EnVision+System-HRP (Dako). For each tumor, the HRP positive cells were counted per low-power (10X objective) field. Each tumor was evaluated with at least six fields in two separate sections.

### Human CD14^+^ monocyte plus T cell co-cultures

PMBCs from healthy donors (n=5) (LifeSource; Evanston, IL) were collected and CD14^+^ monocytes were purified via AutoMacs Magnetic Bead cell separation technology. The CD14-fractions were collected, and CD4^+^ and CD8^+^ T cells purified via AutoMacs Magnetic Bead cell separation technology. The autologous CD4^+^ and CD8+ T cells were cryopreserved for use in the monocyte/T cell co-cultures. The CD14^+^ monocytes were cultured in the presence of medium alone, recombinant human IL-6, recombinant human IL-10, or recombinant human IL-6 plus recombinant human IL-10 (20ng/ml) at 5×10^6^ cells/well in 6-well plates at a final volume of 2ml in cRPMI w/ 10% FCS for 3 days. On day 3 of culture, the non-adherent cells were removed, and the adherent monocytes collected following incubation in presence of 1ml of 1xDPBS containing 2mM EDTA, and incubate the plates at 37°C for 15min. The cryopreserved CD4^+^ and CD8^+^ T cells were thawed, washed, and stained with CFSE. For the macrophage/T cell co-cultures, the cells were cultured in flat-bottomed 96-well plates. The wells were coated with anti-CD3 (0.5μg/ml; 100μl/well; coating wells for 2 hours at 37°C; wells washed 3x with 1x DPBS), and the appropriate amount of Control Ab or anti-B7-H4 was added. For the assessment of T cells proliferation, the T cells were collected, washed in PBS, stained with LIVE/DEAD^®^ Fixable Aqua Dead Cell Stain (Life Technologies; Grand Island, NY), blocked with anti-CD16/32 (ThermoFisher Scientific; San Jose, CA), and then stained with the indicated antibodies - anti-CD4 (clone OKT4) or anti-CD8 (clone 3B5). Results are expressed as the mean percentage of proliferated CD4^+^ and CD8^+^ T cells.. For cytokine analysis replicate wells were harvested on day +3 of co-culture and cytokine secretion determined via multiplex Luminex LiquiChip (Millipore).

### Mouse lymph node cultures

Lymph nodes were collected from naïve SJL/J mice, and cells made into a single cell suspensions. CFSE-labeled total lymph node cells (5×10^5^ cells/well) were cultured in the presence of anti-CD3 (1μg/ml), plus Control Ig, hB7-H4Ig, anti-B7-H4, or hB7-H4Ig plus anti-B7-H4 (5μg/ml). The cells were cultured for 5 days. For the assessment of T cells proliferation, the T cells were collected, washed in PBS, stained with LIVE/DEAD^®^ Fixable Aqua Dead Cell Stain (Life Technologies; Grand Island, NY), blocked with anti-CD16/32 (ThermoFisher Scientific; San Jose, CA), and then stained with the indicated antibodies - anti-CD4 (clone RM4-5). Replicate wells were pulsed with 1μCi of tritiated thymidine at 24 h, and the cultures were harvested on day +5.

PLP_139-151_/CFA Induced EAE and DTH. 6-7 week-old female SJL/J mice were immunized s.c. with 100μl of an emulsion containing 200μg of *M. tuberculosis* H37Ra (BD Biosciences; San Jose, CA) and 50μg of PLP_139–151_ distributed over three sites on the flank. Mice were treated with Control Ig, hB7-H4Ig, and anti-B7-H4 beginning at the time of priming. Mice were treated with 100μg per dose injections 3x per week for 2 weeks via i.p. injection. Individual animals were observed at the indicated time points and clinical scores assessed in a blinded fashion on a 0–5 scale: 0, no abnormality; 1, limp tail; 2, limp tail and hind limb weakness; 3, hind limb paralysis; 4, hind limb paralysis and forelimb weakness; and 5, moribund. The data are reported as the mean daily clinical score. On day +9 post priming, mice were assayed for delayed type hypersensitivity (DTH). Mice were anaesthetized by inhalation of isoflurane and the thickness of both ears was measured using a dial thickness gauge. 10μg of PLP_139-151_ (negative control) and OVA protein in 10μl of PBS was injected into the left and right ear, respectively. The increase in ear thickness was determined after 24 h.

### Flow Cytometry

For cell analysis spleens were dissociated into a single cell suspensions and RBCs lysed. For the bladder leukocytes, Single cell suspensions were prepared by mincing the bladder tissue in 2ml of Accutase (MilliPore) plus 1mg/ml collagenase, and the samples were incubated at 37°C for 30 min. Following the enzyme digestion, the bladder samples were disrupted with a 100μm cell strainer, the cell strainer washed 2x with 10ml of HBSS+ 5% FCS, and the cells pelleted. The cells were washed in PBS, stained with LIVE/DEAD^®^ Fixable Aqua Dead Cell Stain (Life Technologies; Grand Island, NY), blocked with anti-CD16/32 (ThermoFisher Scientific), and then stained with the indicated antibodies. 10^6^ viable cells were analyzed per individual sample using a BD LSRFortessa (BD Bioscience), and the data analyzed using FloJo Version 9.5.2 software (Tree Star, Inc.; Ashland, OR).

| CD4+ T cells | Color | CD8+ T cells | Color | CD4/CD8 | Color | Monocytes | Color |
| --- | --- | --- | --- | --- | --- | --- | --- |
| VID | Live/Dead Blue Indo-violet | VID | Live/Dead Blue Indo-violet | VID | Live/Dead Blue Indo-violet | VID | Live/Dead Blue Indo-violet |
| CD45 (clone 30-F11) | V500 | CD45 (clone 30-F11) | V500 | CD45 (clone 30-F11) | V500 | CD45 (clone 30-F11) | V500 |
| CD3 (clone 145-2C11) | BV650 | CD3 (clone 145-2C11) | BV650 | CD3 (clone 145-2C11) | BV650 | CD11b (clone M1/70) | BV650 |
| CD4 (clone RM4-5) | PB | CD8 (clone 53-6.7) | PB | CD4 (clone RM4-5) | PB | Ly6C (clone AL-21) | Alexa Fluor 700 |
| CD25 (clone PC61) | APC/Cy7 | CD25 (clone PC61) | APC/Cy7 | CD8 (clone 53-6.7) | APC/Cy7 | Ly6G (clone 1A8) | APC/Cy7 |
| CD44 (clone IM7) | BV605 | CD44 (clone IM7) | BV605 | Nrp-1 (polyclonal) | APC | CD80 (clone 16-10A1) | PB |
| FoxP3 (clone FJK-16s) | Alexa Fluor 700 | PD-1 (clone RMP1-30) | PE-Cy7 | Sema3a (mouse and human clone #215) | FITC | CD86 (clone GL1) | APC |
| PD-1 (clone RMP1-30) | PE-Cy7 |  |  |  |  | PD-L1 (clone 10F.9G2) | PE/Cy7 |
|  |  |  |  |  |  | B7-H4 (clone MIH29) | PE |

### Bioinformatics

Clinical, mRNA seq, and mutation data from The Cancer Genome Atlas (TCGA) bladder urothelial carcinoma dataset was downloaded from the Broad Institute Genome Data Analysis Center (GDAC) using TCGA data version 2016_01_28 (http://gdac.broadinstitute.org) (26,27). Analysis was performed with R v3.5.1 utilizing Bioconductor v3.9. RNA-seq mRNA expression levels are presented as RNA-seq by expectation-maximization (RSEM) values, and z-score >2 was used to define overexpression (28). Overall survival outcomes were calculated using the log-rank test and R packages survival v2.43-3 and survminer v0.4.3. Pan-cancer expression profile of B7-H4 (VTCN1) was extracted from the Broad GDAC Firehose v1.1.38 (http://firebrowse.org/). Spearman correlations between B7-H4, PD-1, PD-L1, and CTLA-4 expression and mutation, neoantigen, and APOBEC enrichment was performed using ggplot2 v3.1.0. Molecular subtypes (luminal, luminal-papillary, luminal-infiltrated, basal-squamous, and neuronal) were assigned as previously described (29). Comparison of expression between subtypes was performed using Kruskal-Wallis ANOVA. Immune signatures were calculated as previously described (30–32). Hierarchical clustering of immune signatures in among molecular subtypes was performed with multiClust v1.4.0 and gplots v3.0.1 (33). Correlation heatmap of immune signatures and B7-H4 expression was created using ggplot2 v3.1.0.

For murine tumors, following quality control and adapter trimming, reads were mapped to mm10 genome using STAR (34), and then counted using HTSeq (35). Gene counts were converted to CPM and further analyzed in R using the Bioconductor packages limma and edgeR. Limma was used to generate MDS plots and call differentially expressed (DE) genes. Top DE genes were visualized by Volcano plots, and used for Pathway Analysis that was conducted using the Bioconductor packages TopGO and ReactomePA (36). The most significant pathways (reactomeDB terms) enriched in DE genes were shown by horizontal barplots, where color intensity tracked with p-values (computed by Fisher Exact test).

### Statistical Analysis

Comparisons of the percentage of animals showing clinical disease were analyzed by X^2^ using Fisher’s exact probability, and two-way ANOVA with a Bonferroni post-test was used to determine statistical differences between mean clinical disease scores. Single comparisons of two means were analyzed by Student’s t-test.

## Author Contribution

JRP-designing research studies, conducting experiments, acquiring data, analyzing data, and writing the manuscript, editing manuscript

APG-designing research studies, conducting experiments, acquiring data, analyzing data, and writing the manuscript, editing manuscript

DB designing research studies, conducting experiments, acquiring data, analyzing data, and writing the manuscript, editing manuscript

DF-designing research studies, conducting experiments, acquiring data, analyzing data, and writing the manuscript, editing manuscript

YY designing research studies, conducting experiments, acquiring data, analyzing data, editing manuscript

VE-designing research studies, conducting experiments, acquiring data, analyzing data, editing the manuscript.

SS designing research studies, conducting experiments, acquiring data, analyzing data, providing reagents, and writing the manuscript.

EC designing research studies, conducting experiments, acquiring data, analyzing data, providing reagents, and writing the manuscript.

PR-designing research studies, conducting experiments, acquiring data, analyzing data, writing and editing the manuscript.

SDM-designing research studies, conducting experiments, acquiring data, analyzing data, writing and editing the manuscript.

JJM-designing research studies, conducting experiments, acquiring data, analyzing data, writing and editing the manuscript.

## Acknowledgements

JJM and SDM were supported by a Translational Bridge Grant from the Robert H. Lurie Cancer Center Funded by the John P. Hanson Foundation.

