## Supplemental Figures for "Antibody Targeting of B7-H4 Enhances the Immune Response in Urothelial Carcinoma"

Supplemental Figure 1.

A

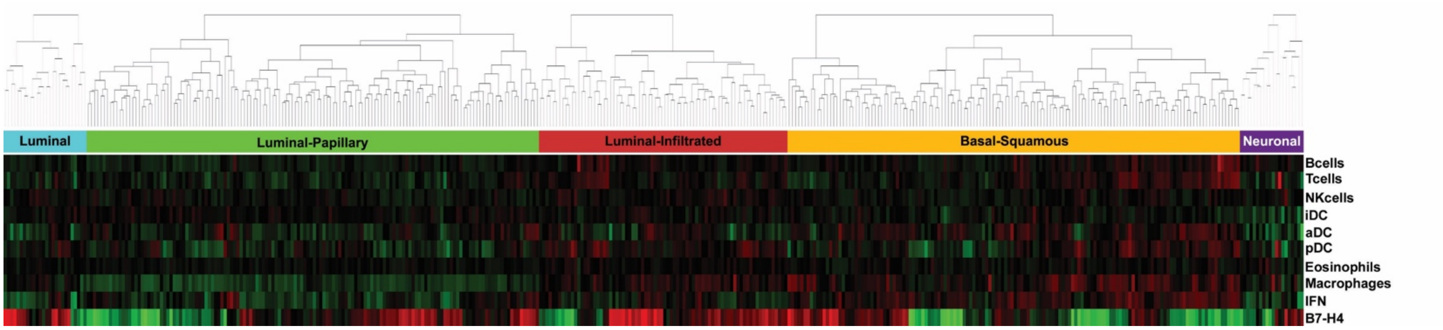


B

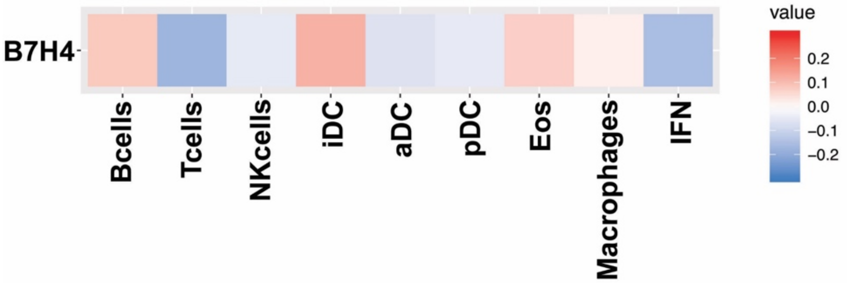


Supplemental Figure 1. Association of *B7-H4* (*VTCN1*) expression with immune signatures in The Cancer Genome Atlas (TCGA) bladder urothelial carcinoma dataset. (A) Hierarchical clustering of immune signatures in luminal, luminal-papillary, luminal-infiltrated, basal-squamous, and neuronal molecular subtypes. (B) Correlation matrix of B7-H4 expression with immune signatures. Value in legend represents Spearman's rho.

NK cells, natural killer; iDC, immature dendritic cell; aDC, activated dendritic cell; pDC, plasmacytoid dendritic cell; IFN, interferon

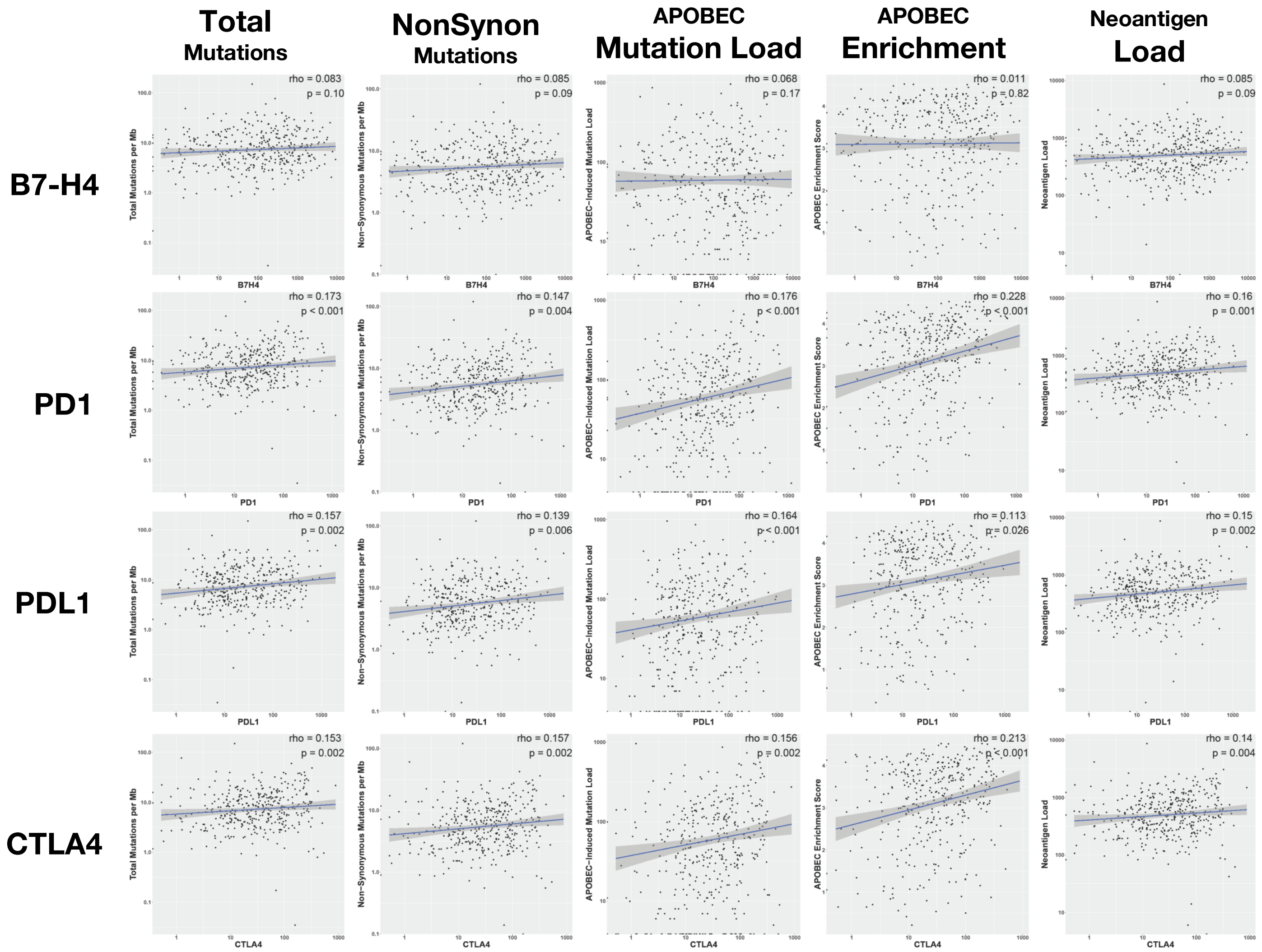


Supplemental Figure 2. Spearman correlation of immune checkpoint expression (RSEM) with mutations and neoantigens in The Cancer Genome Atlas (TCGA) bladder urothelial carcinoma dataset. Rows represent immune checkpoints B7-H4 (VTCN1), PD1 (PDCD1), PD-L1 (CD274), and CTLA4. Columns represent total mutations per Mb, non-synonymous mutations per Mb, APOBEC-induced mutation load, APOBEC-enrichment score, and neoantigen load.

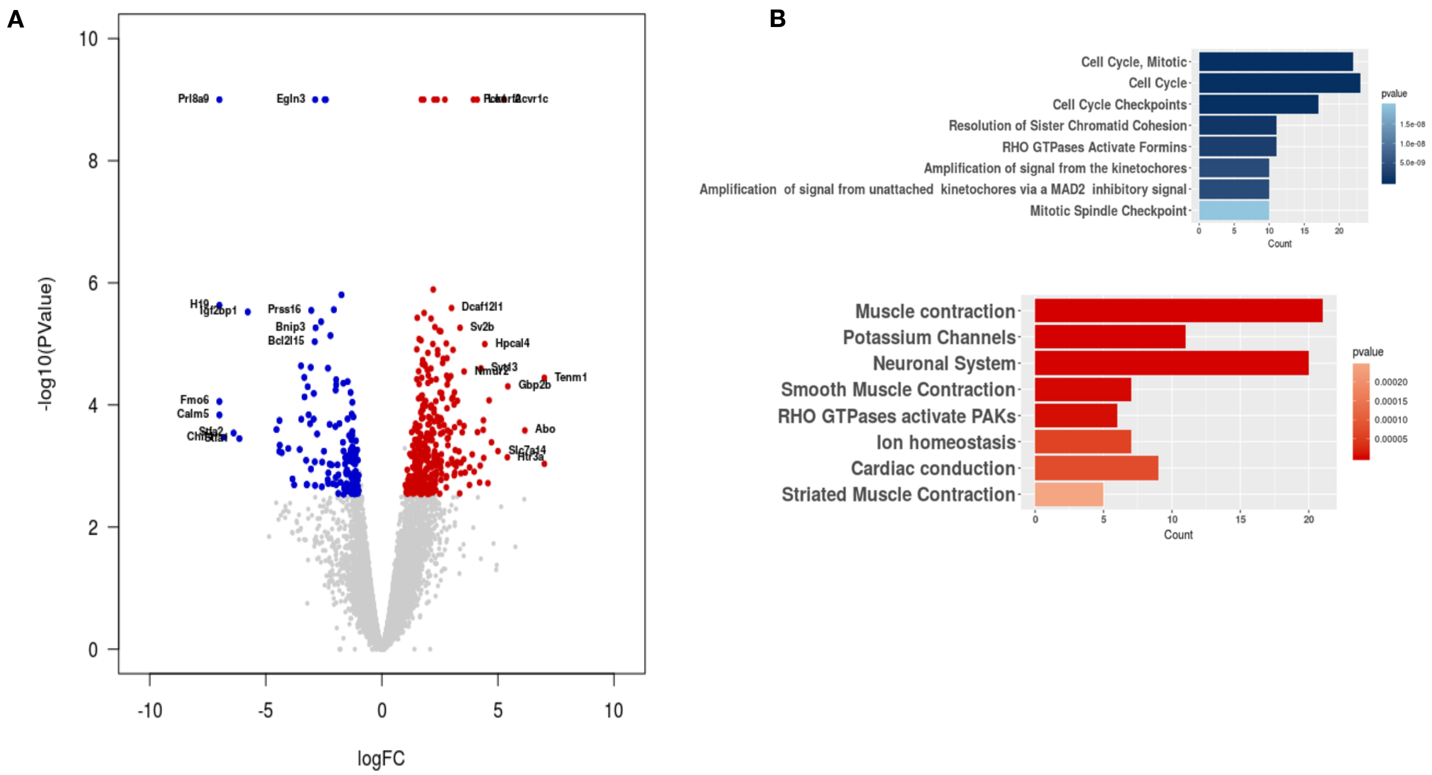


Supplemental Figure 3. Comparison of gene plot of tumors treated with anti-B7H4 with a pathologic response (red) compared to tumors treated with IgG (blue). B) Gene-ontology pathways of reduced (blue) and upregulated (red) gene expression pathways.
